# Dynamic Transcriptome, Accessible Genome and PGR Cistrome Profiles in the Human Myometrium

**DOI:** 10.1101/806950

**Authors:** San-Pin Wu, Matthew L. Anderson, Tianyuan Wang, Lecong Zhou, Olivia M. Emery, Xilong Li, Francesco J. DeMayo

## Abstract

The myometrium undergoes structural and functional remodeling during pregnancy. We hypothesize that myometrial genomic elements alter correspondingly in preparation for parturition. Human myometrial tissues from nonpregnant (NP) and term pregnant (TP) human subjects were examined by RNAseq, ATACseq and PGR ChIPseq assays to profile transcriptome, assessible genome and PGR occupancy. NP and TP specimens exhibit 2890 differentially expressed genes, reflecting an increase of metabolic, inflammatory and PDGF signaling, among others, in adaptation to pregnancy. At the epigenome level, patterns of accessible genome change between NP and TP myometrium, leading to altered enrichment of binding motifs for hormone and muscle regulators such as the progesterone receptor (PGR), Krüppel-like factors and MEF2A transcription factors. PGR genome occupancy exhibits a significant difference between the two stages of the myometrium, concomitant with distinct transcriptomic profiles including genes such as *ENO1*, *LHDA*, and *PLCL1* in the glycolytic and calcium signaling pathways. Over-representation of SRF, MYOD and STAT binding motifs in PGR occupying sites further suggests interactions between PGR and major muscle regulators for myometrial gene expression. In conclusion, changes in accessible genome and PGR occupancy are part of the myometrial remodeling process and may serve as mechanisms to formulate the state-specific transcriptome profiles.

## Introduction

Myometrium is the muscular component of the uterus consisting of primarily smooth muscle and vascular cells. The main function of the myometrium is to maintain structural integrity of uteri during pregnancy and to provide contraction force for parturition. Myometrial smooth muscle cells increase in number and size as well as change from a synthetic to a contractile state during gestation [1, 2]. These cells need to remain quiescence before parturition and produce phasic contractions at the laboring stage [3]. After parturition, the involution process reduces the size of uterus to the nonpregnant state [4, 5]. Disorders associated with the myometrium may result in preterm birth, dystocia, adenomyosis and leiomyoma, which impose serious burdens on maternal and child health.

Steroid hormones progesterone, estrogen and glucocorticoid have been indicated for regulating gene expression and cell proliferation of myometrial smooth muscle [6–10]. Estrogen modulates contraction-associated genes and induces hyperplasia of myometrial smooth muscle cells at early gestation by activating the PI3K/mTOR pathway and repressing myostatin expression [7, 8, 11, 12]. Estrogen also supports the expansion of myometrial progenitor-like cells [13] and works together with progesterone to promote proliferation of leiomyoma side-population cells through regulation of the paracrine WNT signaling from the mature myometrial cells. [14]. On the other hand, progesterone signaling downregulates expression of genes that are associated with cell proliferation in cultured human myometrial cells and has a negative effect on vascular smooth muscle proliferation [9, 15, 16]. In contrast, glucocorticoids inhibits proliferation of uterine leiomyoma cells, vascular smooth muscle cells and airway smooth muscle cells but has no impact on the cell cycle progression in cultured uterine smooth muscle cells[6, 17, 18]. The role of progesterone and glucocorticoid in the early gestation myometrium remains unknown.

Progesterone is also an FDA-approved tocolytic agent to prevent premature parturition [19]. Progesterone signaling is primarily mediated by the progesterone receptor (PGR) and is sensitive to environmental endocrine disruptor [16, 20]. Mechanisms that control progesterone signaling include metabolism of progesterone, transcription regulation and post translational modifications of PGR and interaction between PGR and coregulators [21–25]. Before parturition, myometrial progesterone signaling suppresses inflammatory and contractile activities via the ZEB1/ZEB2-miR-200-STAT5b pathway, the PGR-AP-1 containing transcription repressive complex and cAMP-DUSP1 dependent regulation of PGR phosphorylation [23, 26, 27]. At the transition to the laboring stage, emerging reports suggest that the elevated ratio of PGR isoform A verse B could turn the effect of progesterone signaling from repressive to inductive on transcription of labor genes such as GJA1 and further promotes gap junction coupling in the myometrium [27–29]. These findings demonstrate the versatility of the progesterone signaling in the myometrial biology manifested by the activity of its cognate receptor PGR.

This study focuses on the physiological change between nonpregnant and term pregnant myometrium. The profiles of transcriptome, open chromatin regions and PGR cistrome in human myometrial specimens at both stages are documented to identify genetic pathways and potential underlying mechanisms that may be responsible for the structural and functional adaptation to pregnancy. The common and distinct molecular features of myometrium at these two stages not only provide clinical relevance to known pathways but also suggest novel mechanisms that may contribute to myometrial remodeling in support of pregnancy.

## Materials and Methods

### Human Myometrial Specimens

Permission to work with human tissue specimens for all experiments was obtained from the Institutional Review Board for Baylor College of Medicine (BCM H-22119, Approved 5/7/2008). All specimens were obtained from the Gynecologic Tissue Repository for BCM, which has previously been granted permission to routinely collect specimens of female reproductive tract tissues under a separate IRB protocol (H-33461). Written informed consent is available for all specimens utilized to perform the work described.

For the purposes of these experiments, specimens of myometrium were collected from non-laboring gravid women at term (TP; 38-42 weeks estimated gestational age) undergoing cesarean section for routine clinical indications. All myometrial specimens from gravid women were collected from the margins of the hysterotomy following delivery of the fetus. No specimens were collected from women who had undergone pitocin-induced augmentation of labor within 6 hours of surgery or who had previously received 17-hydroxyprogesterone for the treatment of pre-term labor. In addition, no specimens were collected from women whose pregnancy had been complicated by other medical conditions (including but not limited to gestational diabetes >IB, chorioamnionitis, pregnancy-induced hypertension, preeclampsia, HELLP syndrome or its variants). Specimens of non-gravid myometrium (NP) were obtained from healthy, pre-menopausal women undergoing routine, clinically-indicated gynecologic surgery. All non-gravid myometrial specimens were collected from healthy myometrium at the proliferative phase of the menstrual cycle and at sites that excluded both the uterine serosa and endometrium and were at least 2 cm from the nearest leiomyoma. Any specimens from women using progestins prior to their surgery were excluded from analysis.

After washing each specimen briefly in phosphate buffered saline (PBS), pieces of myometrium were either flash frozen (ChIPSeq) or preserved by plunging them into ice-cold RNAlater (RNASeq) with warm ischemia time less than 30 minutes. All specimens were subsequently stored at <−80°C until use. Assays for each individual sample are listed in Table S1.

### RNA Extraction

Myometrial specimens were homogenized by the bead Mill 24 homogenizer (15-340-163, Fisher Scientific, Waltham, MA) in Bead Mill Tubes (15-340-154 Fisher Scientific, Waltham, MA) with 1 mL Trizol (15596026, Thermo Fisher Scientific, Waltham, MA). Tissue debris was pelleted and removed by centrifugation at 12000 xg for 10 minutes at 4°C. After adding 200 uL 1-Bromo-3-chloropropane, samples were manually shake for 20 seconds followed by incubation at room temperature for 3 minutes. Phase separation was conducted by centrifugation at 12000 xg for 18 minutes at 4°C. The RNA containing aqueous layer was retained and subsequently mixed with 500 uL 200 proof ethanol. The mixture was then passed through the RNA binding column of RNeasy Mini Kit (74104, Qiagen, Germantown, MD), followed by washing and elution steps described in the manufacture’s handbook.

### RNAseq

Three NP and three TP myometrial specimens were subject to the RNAseq assay. The libraries prepared from myometrial specimens were sequenced with approximately 50 million, 101bp paired-end reads per sample. The raw reads were subsequently processed by filtering with average quality scores greater than 20. Then the reads were aligned to hg19 using TopHat version 2.0.4 [30]. Expression values of RNAseq were expressed as FPKM (fragments per kilobase of exon per million fragments) values. Differential expression was calculated using Cufflink version 2.2.1 [31]. Differentially expressed genes are defined as absolute fold change ≥ 1.4, adjusted p-value < 0.05 and having FPKM ≥ 1 in at least one of the samples. Functional analysis of gene lists was performed using the Ingenuity Pathway Analysis (IPA, www.ingenuity.com) and The Database for Annotation, Visualization and Integrated Discovery (DAVID) v6.8 [32, 33].

### ATACseq

Three nonpregnant (NP) and 3 term pregnant (TP) myometrial specimens were sent to Active Motif (Carlsbad, CA) for the library preparation service (catalog number 25079). The libraries were generated with the Illumina Nextera DNA Library Prep kit and were sequenced by Active Motif and NIEHS Sequencing core by Illumina HiSeq 2000 and NextSeq 500 systems with 42bp paired-end reads on each sample. The raw reads were initially processed by trimming adaptors and filtering with average quality scores greater than 20. The reads passing the initial processing were aligned to the human reference genome (hg19) using bowtie version 1.1.2 [34]. After removing the reads mapped to mitochondria DNA, the uniquely mapped deduplicated reads in each sample were normalized by down-sampling to 91.5M single reads. Only the first 9bp of each read were used for downstream analyses. The open chromatin regions (OCRs) were first identified by MACS2 [35] with a cutoff of adjusted p-value 0.0001, followed by merging genomic intervals within 100bp of each other.

Motif analysis and peak annotation to nearby gene within 25Kb were performed by the Hypergeometric Optimization of Motif EnRichment (HOMER) motif discovery tool [36]. The bivariate genomic footprinting (BaGFoot) algorithm [37] was used to identify binding motifs exhibiting change of chromatin accessibility including footprint depth and flanking accessibility between 2 myometrial stages with default settings.

### ChIPseq

Two NP and two TP myometrial specimens were subject to the ChIPseq assay. Chromatin immunoprecipitation and library preparation and sequencing were performed by Active Motif (catalog number 25006 and 25046, Carlsbad, CA) using an anti-PGR antibody (sc-7208, Santa Cruz Biotechnology, Dallas, TX) and a custom Illumina library with the standard Illumina PE adaptors [38]. The Illumina HiSeq 2000 system was utilized for next generation sequencing. The 50-nt single-end sequence reads identified were mapped to genome assembly hg19 using Bowtie version 1.1.2 with default settings. Only reads that passed Illumina’s purity filter and of a mean quality score of 20 or higher, aligned with no more than 2 mismatches, and mapped uniquely to the genome were used in the subsequent analysis. In addition, unless stated otherwise, duplicate reads were removed using the MarkDuplicates tool available in picard-tools-1.119. The retained read alignments were sorted by coordinates, extended to 300 bases, and then randomly downsampled to 9 million intervals per sample with custom scripts. MACS (version 2.1.1.20160309) was employed to perform peak calling with FDR cutoff at 0.0001, using a human myometrium input as control. HOMER’s “annotatePeaks.pl” and findMotifsGenome.pl” functions with default settings were used for peak annotation and motif enrichment analysis of given peak ranges [36]. BEDTools version 2.27.1 was chosen for merging and intersecting peaks in overlapping analysis, with the default settings [39].

### ChIP-qPCR

Two NP and two TP samples independent of the specimens used in ChIPseq were subject to ChIP-qPCR assay. ChIP-qPCR was performed by Active Motif (Carlsbad, CA) using an anti-PGR antibody (sc-7208, Santa Cruz Biotechnology, Dallas, TX). Primer sequences for the ChIP-qPCR assay are ENO1_cF: AGCCCTTCCCCAATCATTAC, ENO1_cR: TACGTTCACCTCGGTGTCTG, LDHA_cF: GTGCATTCCCGGTACGGTAG, LDHA_cR: AGCAGAACCAGAGGCAGTTG, PLCL1_cF: GGAGAATGCGGGTCATCTTG, PLCL1_cR: CAGGAAACAAGCAGGAGTAGG and Untr12 (Catalog #71001, Active Motif, Carlsbad, CA). PGR binding events were calculated according to an equation published in the user’s manual of Active Motif catalog number 12970399.

### Quantitative RT-PCR

Six NP and nine TP samples independent of the specimens used in RNAseq were subject to the quantitative RT-PCR assay. cDNA was generated by the Transcriptor First Strand cDNA Synthesis Kit (04379012001, Roche Life Science, Penzberg, Germany) in accordance to the manufacture’s manual. PCR was conducted by using SsoAdvanced Universal SYBR Green Supermix (1725270, Bio-Rad, Hercules, CA) or SsoAdvanced Universal Probes Supermix (1725280, Bio-Rad, Hercules, CA) on the CFX Connect Real-Time PCR Detection System (1855201, Bio-Rad, Hercules, CA) based on manufacture’s instruction.

Primers used in this study include OXTR_F: GGGGAGTCAACTTTAGGTTCG, OXTR_R: TTCCTCGGGATGTTCAGC, GJA1_F: GCCTGAACTTGCCTTTTCAT, GJA1_R: CTCCAGTCACCCATGTTGC, ZEB1_F: CCTAAAAGAGCACTTAAGAATTCACAG, ZEB1_R: CATTTCTTACTGCTTATGTGTGAGC, PLCL1_F: CAGGAAAAGATTGTACAGTGTCAGA, PLCL1_R: TTTGCCCCCAAATTATGAAG, SLC2A1_F: GGTTGTGCCATACTCATGACC, SLC2A1_R: CAGATAGGACATCCAGGGTAGC, HK2_F: GCTGAAGGAAGCGATCCA, HK2_R: GTGTCGTTCACCACAGCAAC, ALDOB_F: GGCAAGGCTGCAAACAAG, ALDOB_R: CCCGTGTGAACATACTGTCCT, PGK1_F: CAGCTGCTGGGTCTGTCAT, PGK1_R: GCTGGCTCGGCTTTAACC, ENO1_F: TCCCAACATCCTGGAGAATAA, ENO1_R: ATGCCGATGACCACCTTATC, PKM2_F: ACGTGGATGATGGGCTTATT, PKM2_R: CCAAGGAGCCACCATTTTC, LDHA_F: GTCCTTGGGGAACATGGAG and LDHA_R: TTCAGAGAGACACCAGCAACA. The 18S rRNA probe is from ThermoFisher Scientific (4319413E, ThermoFisher Scientific, Waltham, MA). All the RT-PCR results were normalized to the expression levels of the internal control 18S rRNA.

### Statistical Analysis

Two tailed Student’s t-test with equal variance was used for the quantitative qPCR assays.

### High Content Data Deposit

The RNAseq, ChIPseq and ATACseq raw data is under the SuperSeries GSE137552 in the Gene Expression Omnibus of the National Center for Biotechnology Information at the National Library of Medicine.

## Result

### Adaptive Myometrial Transcriptome for Pregnancy

Previous studies show that the expression of parturition associated genes *OXTR* and *GJA1* increase and *ZEB1* and *PLCL1* decrease expression at the transition from quiescence to labor in the term pregnant myometrium [40–43]. However, the expression levels of these genes in the myometrium prior to term remains unclear. After examining 6 nonpregnant (NP) and 9 term pregnant (TP) myometrial specimens, we extend these findings by demonstrating that mRNA levels of *OXTR, GJA1, ZEB1* and *PLCL1* genes all start lower in the nonpregnant myometrium and rise toward term pregnancy (Figure 1). These observations suggest that the myometrial transcriptome is dynamically staged in between the two states of myometrium in preparation for parturition.

**Figure 1.**
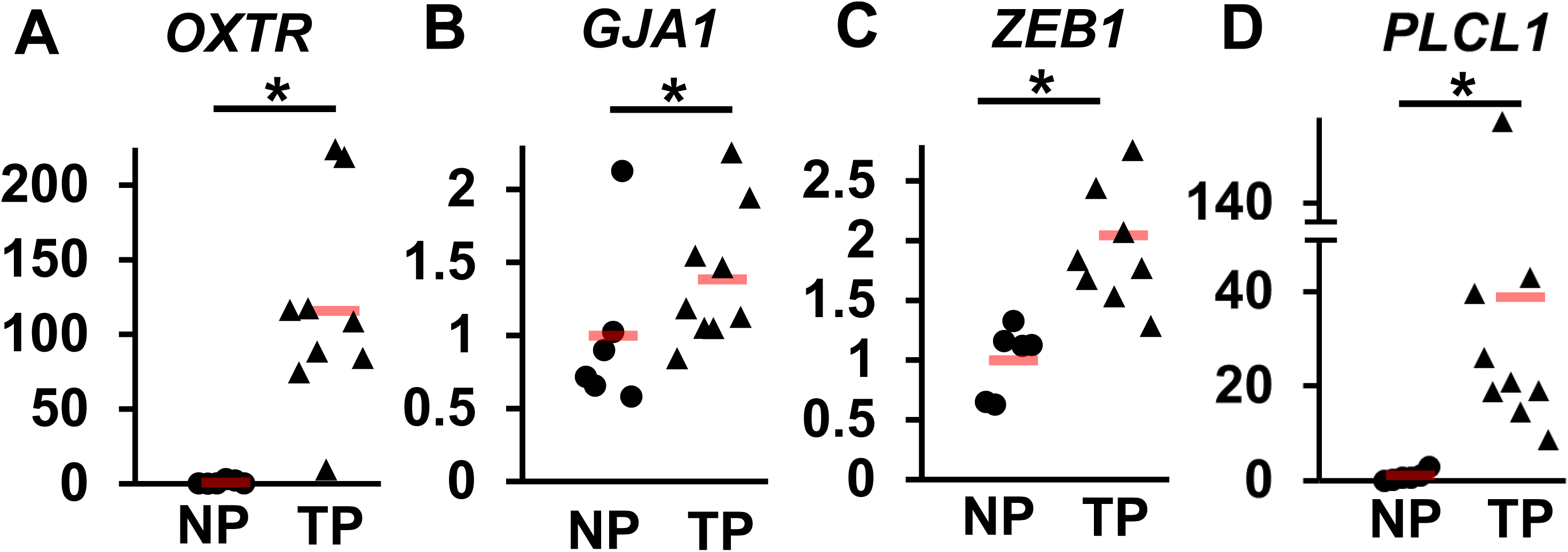
mRNA levels of parturition related genes in nonpregnant and term pregnant myometrium. Relative mRNA levels by qRT-PCR from myometrial tissues of 6 (NP, dots) and 9 (TP, triangles) human subjects. The Y-axis denotes relative mRNA levels. *, p < 0.05 by two-tailed t-test. Orange bars denote the mean of each group.

Genome-wide transcriptome dynamics from the NP to the TP stage was further investigated by RNAseq in a cohort of myometrial specimen independent from those in the aforementioned study of marker genes (Table S1). Using a criteria of average fragments per kilobase of transcript per million mapped reads (FPKM) greater or equal than 1 in at least one of the six specimens, 15907 active genes were found in the myometrium (Table S2). Within each group, the NP and TP myometrium each has 14899 and 15260 active genes, respectively (Table 1 and Table S2). Between these two stages, NP and TP samples share 14612 active genes (Table S2). While 95.7% NP and 93.4% TP expressing genes are present in both stages (Table 1), overall transcriptome profiles exhibited a significant difference as evidenced by the principle component analysis that shows distinct separation between samples of TP and NP stages (Figure 2A). These findings suggest that the pregnancy dependent transcriptome dynamics in the myometrium is primarily a result of change of mRNA levels rather than the composition of expressing genes.

**Figure 2.**
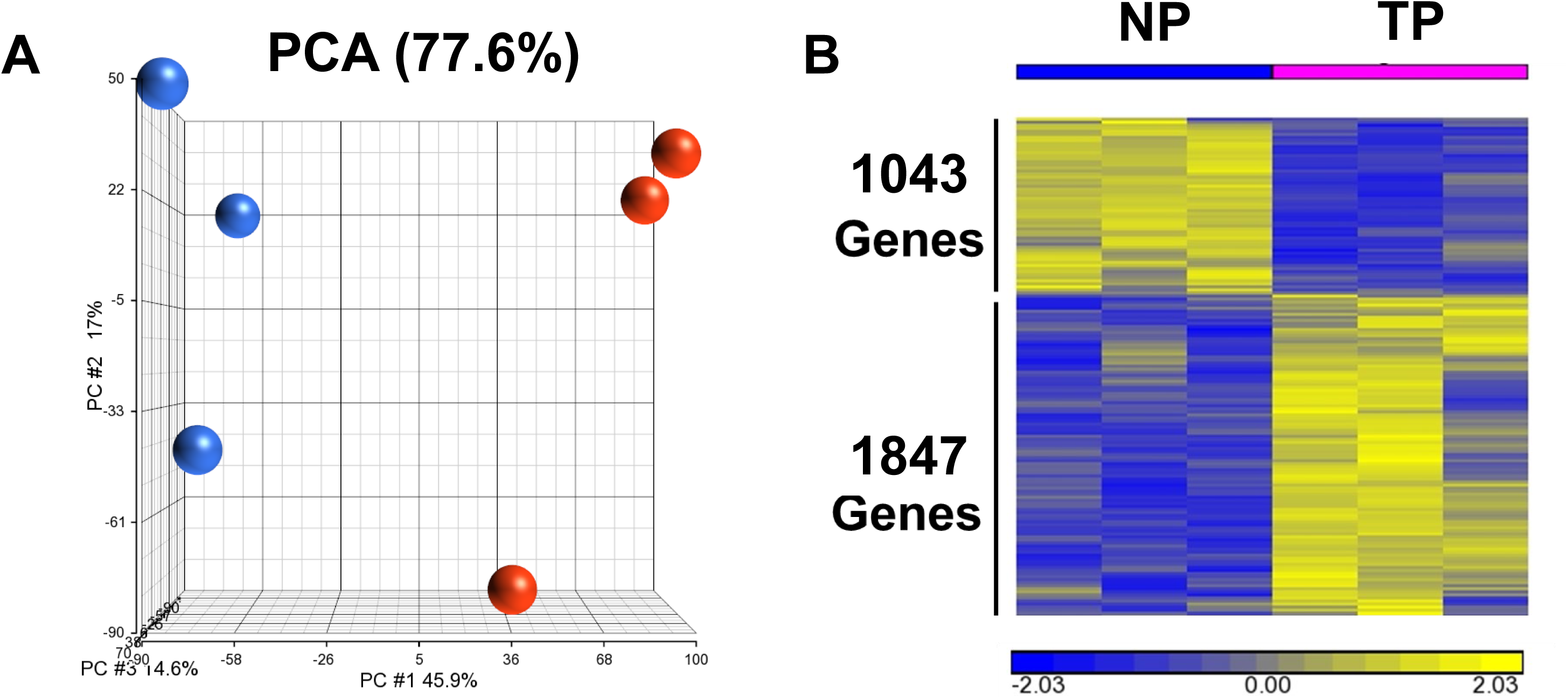
Differences between NP and TP myometrial tissues at the transcriptomic level. (A) Principle Component Analysis of NP (blue) and TP (red) Transcriptome. (B) Differentially expressed genes (DEGs) between groups. Differentially expressed genes are defined as absolute fold change ≥ 1.4, adjusted *p*-value < 0.05 and having FPKM ≥ 1 in at least one of the samples.

**Table 1.**
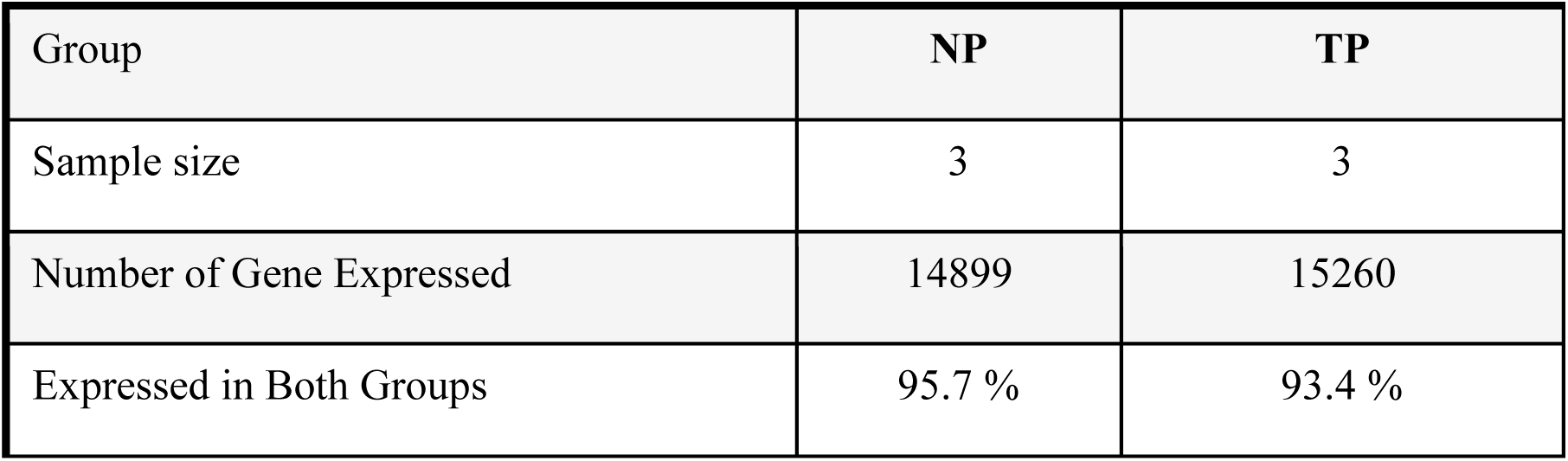
Transcriptome of Human myometrium in Non-pregnant (NP) and Term-pregnant (TP) human subjects. Genes with FPKM ≥ 1 in at least one of the specimens within each group are defined as expressed.

| Group | NP | TP |
| --- | --- | --- |
| Sample size | 3 | 3 |
| Number of Gene Expressed | 14899 | 15260 |
| Expressed in Both Groups | 95.7 % | 93.4 % |

Using the absolute fold change greater or equal to 1.4 and adjusted *p*-value less than 0.05 as the cutoff, 2890 differentially expressed genes (DEGs) were identified between the two stages of myometrium with expression of 1043 and 1847 genes relatively higher in NP and TP tissues, respectively (Figure 2B). Parturition related genes again show increased expression toward term pregnancy in this cohort of specimens with mean fold changes at 156 for *OXTR*, 2.9 for *GJA1*, 1.6 for *ZEB1* and 2.6 for *PLCL1* (Table S2), consistent with previous observations in another cohort of specimens (Figure 1). Gene ontology analysis on the DEGs shows that genes expressed at higher levels at the NP stage are enriched in terms in association with transcription, RNA-binding and histone deacetylase activity among others, reflecting the synthetic state of the NP myometrium. In contrast, terms linked to innate immunity, protease and carbohydrate transport and metabolism are over-represented among genes of higher expression levels at pregnancy (Table 2 and Table S3), demonstrating a molecular profile of the contractile state in favor of inflammation control, extracellular remodeling and increased energy demand as well as in preparation for subsequent parturition. Further examining the 2890 DEGs reveals potential alterations of molecular activities between NP and TP stages, as predicted by the differential expression of downstream target genes of molecules of interest. The Ingenuity Pathway Analysis (IPA) estimates increased activities of myometrial expansion/remodeling associated TGFβ, PDGF, ERK and STAT, inflammation management linked TNF and AP-1, as well as myometrial contraction related p53 signaling, among others (Table 3 and Table S4) [11, 44–49]. Changes of these molecular activities appear to be evolutionarily conserved, as evidenced by a similar observation that compares transcriptomic profiles of the mouse myometrium between 18.5 days post coitus and virgin (GSE17021, Table 3 and Table S5). Notably, majority of the subset of DEGs that match IPA curated glucose downstream exhibits changes of gene expression consistent with the response to glucose treatment (Table 3, Table S4 and Table S5), which suggests relatively higher glucose utilization at the TP stage. This result is also in line with the aforementioned gene ontology enrichment finding as shown in Table 2. Moreover, genes that encode enzymes of the glycolysis pathway, including *SLC2A1, ALDOB, PGK1, ENO1, PKM2* and *LDHA*, exhibited increased mRNA levels toward term pregnancy in two independent cohorts of specimens (Figure 3 and Table S2). The HK2 gene, which encodes the hexokinase for glucose phosphorylation, also showed significant higher expression levels in the TP myometrium in the first cohort despite displayed a trend of increase in the second cohort instead (Figure 3). The higher expression in glycolytic pathway genes and glucose downstream targets together suggest an increase of glucose usage in the TP myometrium compared with that in the NP stage. Taking together, these findings illustrate an adaptive myometrial transcriptome in response to the pregnancy and in preparation for parturition at various aspects such as structure reorganization, inflammation and contraction management, and metabolic adaptation.

**Figure 3.**
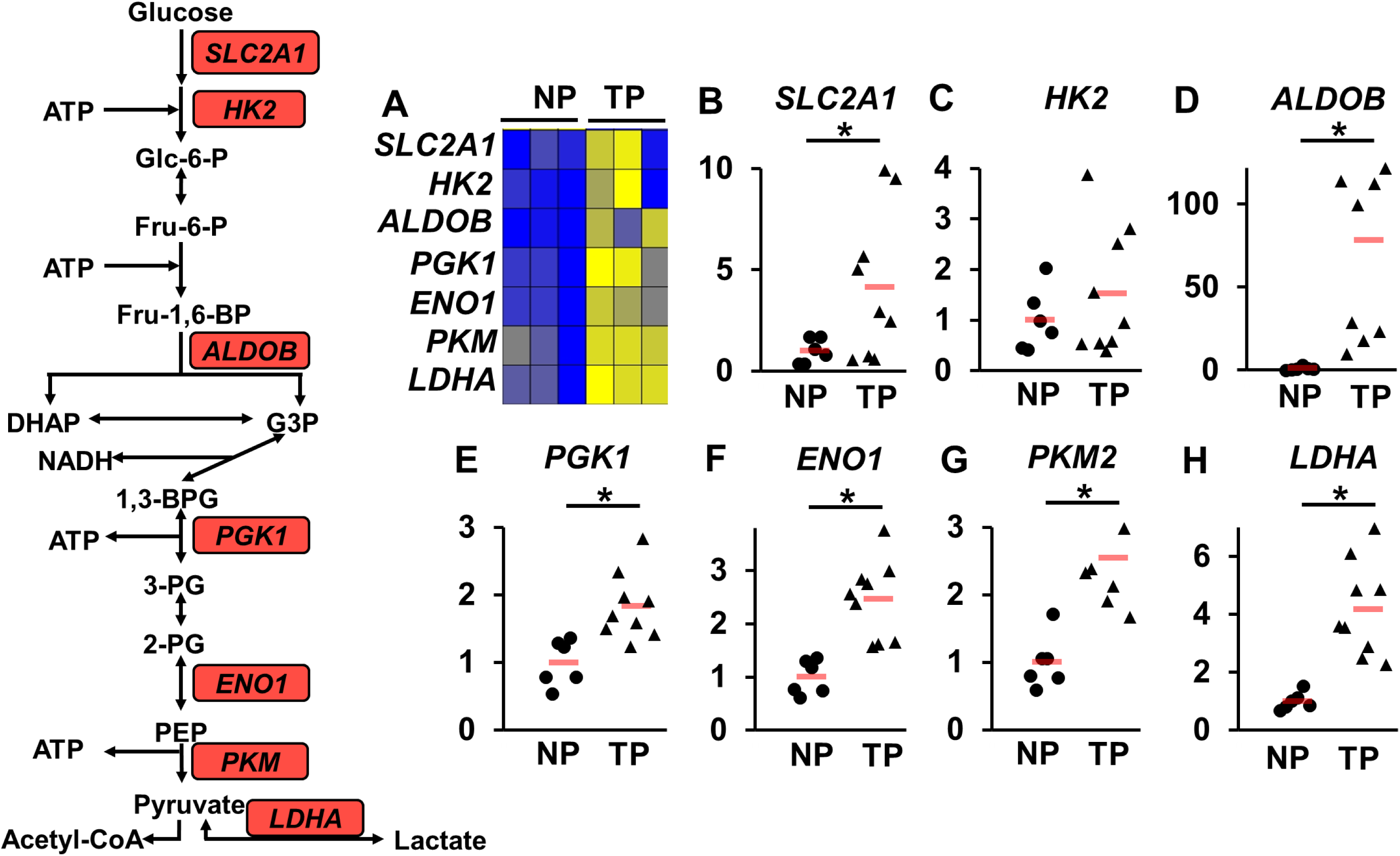
Expression of glycolytic genes. The diagram to the left depicts the glycolysis pathway. (A) RNAseq results. Yellow and blue depict high and low levels of gene expression, respectively. N=3 for NP and 3 for TP. (B-H) Relative mRNA levels of tissues from the same cohort of human subjects as depicted in figure 1. The Y-axis denotes relative mRNA levels. *, *p* < 0.05 by two-tailed t-test. Orange bars denote the mean of each group. N=6 for NP and 9 for TP.

**Table 2.**
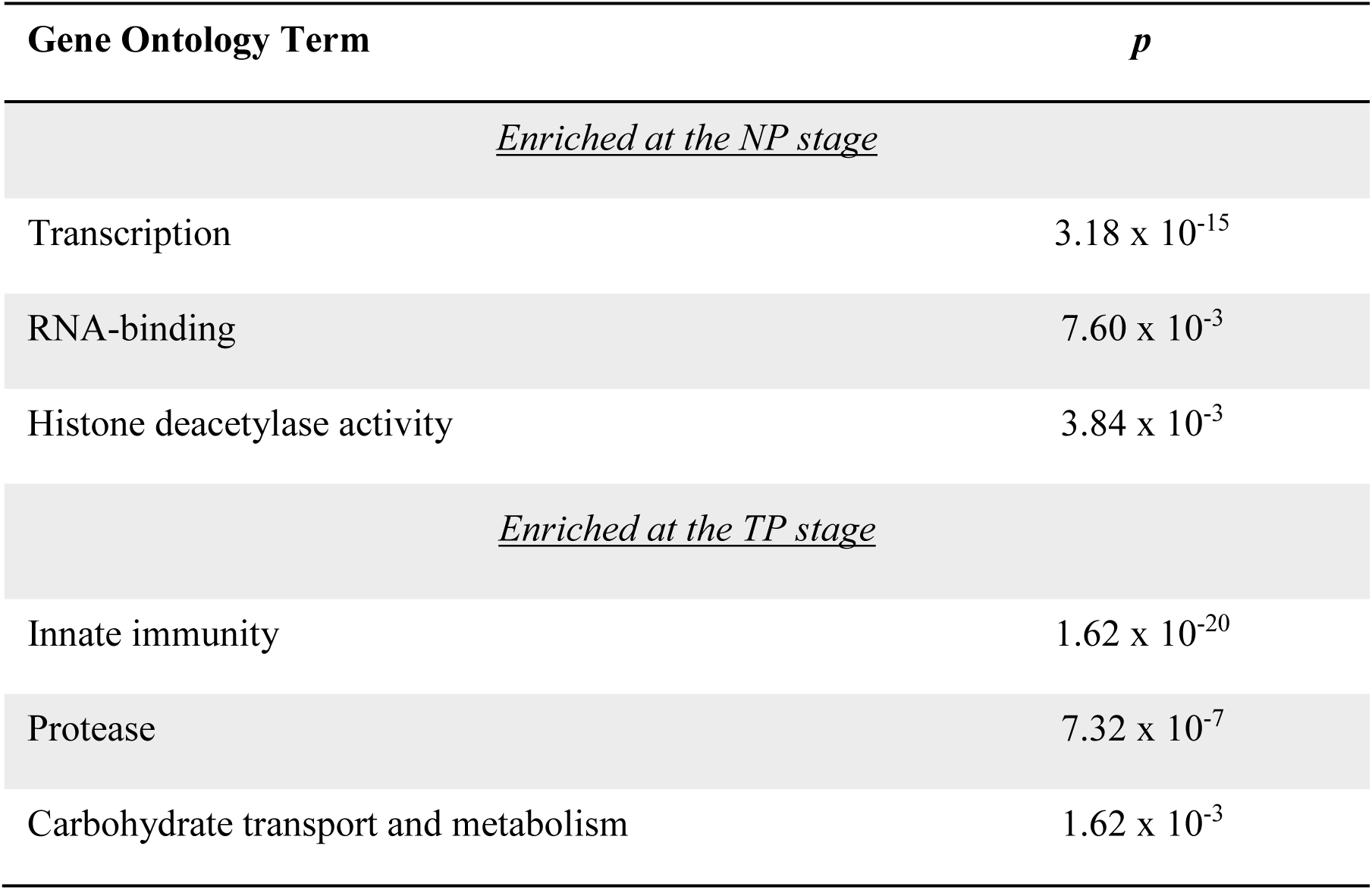
Enriched Gene Ontology terms in differentially expressed genes (DEGs) between NP and TP myometrium.

**Table 3.** Changes of molecular activities from NP to TP predicted by Ingenuity Pathway Analysis (IPA) based on human RNAseq and mouse microarray (GSE17021, virgin vs. 18.5 dpc myometrium) results.

| **Change of Molecular activities** | **Human Activation**<br>**z-score** | **Human Enrichment**<br>***p* value** | **Mouse Activation**<br>**z-score** | **Mouse Enrichment**<br>***p* value** |
| --- | --- | --- | --- | --- |
| TGFβ | 2.40 | 1.05 x 10<sup>-15</sup> | 4.38 | 2.13 x 10<sup>-13</sup> |
| PDGF-BB | 2.82 | 1.67 x 10<sup>-18</sup> | 3.49 | 9.03 x 10<sup>-17</sup> |
| PI3K | 5.04 | 2.38 x 10<sup>-14</sup> | 2.57 | 4.16 x 10<sup>-10</sup> |
| TNF | 8.31 | 2.92 x 10<sup>-73</sup> | 2.36 | 6.75 x 10<sup>-43</sup> |
| AP-1 | 2.41 | 3.66 x 10<sup>-12</sup> | 2.16 | 8.12 x 10<sup>-5</sup> |
| ERK | 3.87 | 3.44 x 10<sup>-12</sup> | 3.11 | 3.85 x 10<sup>-8</sup> |
| p53 | 2.39 | 5.00 x 10<sup>-53</sup> | 3.03 | 4.02 x 10<sup>-54</sup> |
| STAT3 | 2.87 | 5.04 x 10<sup>-27</sup> | 1.95 | 1.97 x 10<sup>-6</sup> |
| D-Glucose | 2.39 | 2.44 x 10<sup>-20</sup> | 2.63 | 9.84 x 10<sup>-18</sup> |

### Accessible Genome Dynamics in the Myometrium

Since the myometrial transcription profiles are adaptive to pregnancy, we next examine the myometrial genome accessibility to check if the profiles of open chromatin regions are also different between NP and TP states. The assay for transposase-accessible chromatin using sequencing (ATACseq) [50] was performed on myometrial specimens of three human subjects of each stages. The NP and TP myometrium each has 45501 and 85777 open chromatin regions (OCRs) identified in at least one of three samples within the group (Figure 4 and Table S6). Among these genomic regions, 9564 (class I) and 50840 (class III) OCRs are exclusively present in the NP and TP states, respectively, whereas 35937 OCRs (class II) can be detected in both states (Figure 4). Within each class of OCRs, HOMER identified enrichment of 128, 82 and 121 transcription factor binding motifs in class I, II and III OCRs, respectively (Table S7). Among these enriched motifs, 45, 3 and 13 motifs show enrichment uniquely present in class I, II and III OCRs, respectively (Table 4). Genes that encode corresponding transcription factors for vast majority of these motifs are found active in human myometrial tissues (Table S2), implicating that interactions of these transcription factors with genomic DNA have a high probability to be affected by change of chromatin structure between the two states of myometrial tissues.

**Figure 4.**
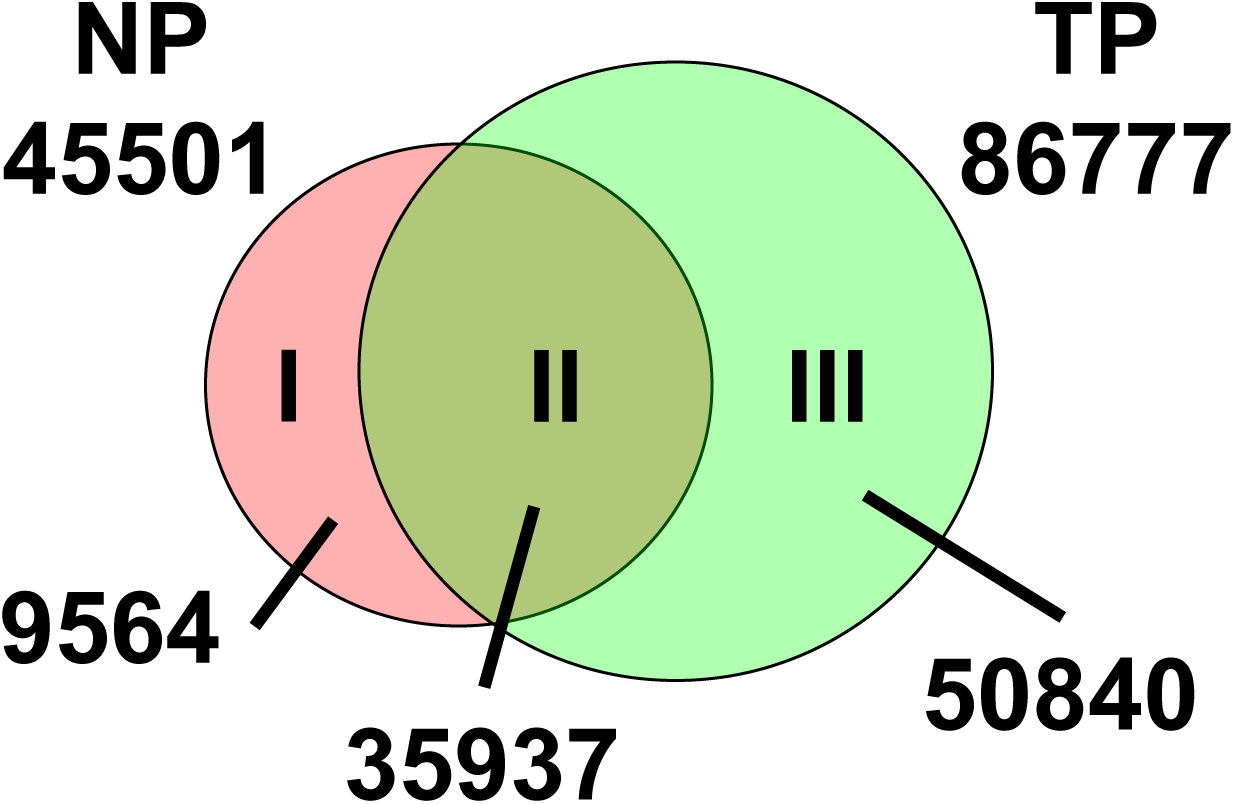
Open chromatin region in the myometrium. Numbers of open chromatin regions (OCRs) identified in 3 NP and 3 TP specimens by ATACseq. OCRs within each group were pooled together. Numbers of shared and unique OCRs between NP and TP samples are also noted.

**Table 4.** Enriched motifs in each class of open chromatin regions.

| **Class** | **Number of Total Enriched Motifs** | **List of Uniquely Enriched Motifs** |
| --- | --- | --- |
| I | 128 | AMYB, Arnt:Ahr, Ascl1, Atoh1, BMYB, c-Myc, E2F1, E2F6, Egr1, Eomes, ERE, Esrrb, Fox:Ebox, FOXA1, FXR, Gfi1b, HNF4a, HOXA2, KLF5, Lhx2, Lhx3, Max, MITF, MYB, MyoD, Nanog, NeuroD1, Nkx6.1, PAX5, Pax8, Pdx1, PPARE, PGR, PRDM1, PRDM9, PU.1-IRF, Rbpj1, RUNX1, RUNX2, RXR, Smad2, SPDEF, TATA-Box, THRa, VDR |
| II | 82 | GFX, GFY, ZBTB33 |
| III | 121 | CEBP, EKLF, Hnf1, HRE, Mef2a, NF1, Nr5a2, Nur77, PAX3, RORgt, RUNX, Tbx20, Usf2 |

Class I and III OCRs that are located within 25 kb of transcription start sites of DEGs are arbitrarily defined as DEG-associated dynamic OCRs (Table 5), rendering these OCRs the potential gene regulatory elements for linked DEGs subject to the control of chromatin structure changes. Among the linked OCRs and DEGs, positive correlation is observed between 2989 OCRs and 1461 DEGs that have elevated gene expression concomitant with increased genome accessibility or decreased gene expression with reduced genome accessibility (Table 5). This observation suggests that these 2989 OCRs could function as putative enhancers for the 1461 associated DEGs. On the other hand, negative correlation is seen between 1660 OCRs and 980 DEGs that exhibit a reduction of gene expression with increased genome accessibility or an increase of gene expression with decreased genome accessibility (Table 5). This inverse correlation implicates a role of these 1660 OCRs as putative repressors for the 980 DEGs. Motif enrichment analysis on the DEG-associated dynamic OCRs further reveals candidate transcription regulators that may control transcription of the associated DEGs and whose actions are possibly affected by change of chromatin structure between the two states of myometrium (Table S8). Among the DNA-binding proteins that may occupy these enriched motifs, estrogen receptor, glucocorticoid receptor, progesterone receptor, MEF2C, NFAT, NFkB, AP-1 and STAT transcription factors have been studied for their roles in the myometrial biology [6, 51–54]; and CTCF and YY1 are known for their functions in regulation of chromatin conformation [55, 56]. The presence of these motifs in the dynamic OCRs suggest that changes in chromatin structure may also have an impact on the actions of these transcription and chromatin conformation regulators in modulation of associated DEGs between the two states of myometrium. Further examining the associated DEGs by gene ontology enrichment analysis found signaling pathways that are potentially subject to chromatin structural changes, including Rho family GTPases, actin cytoskeleton, gap junction, eNOS, Relaxin, PDGF and HIPPO signaling for myometrial remodeling and contraction regulation as well as glycolysis and Adrenomedullin for metabolism and circulation control (Table 6 and Table S9). In summary, these findings identify potential signaling pathways and their putative transcriptional regulators in the human myometrium that may have an additional control mechanism through genome accessibility.

**Table 5.** Association between DEGs and dynamic OCRs. Class I and III open chromatin regions are defined as DEG-associated if they are located within 25kb of the transcription start site of DEGs. Positive correlation is defined if 1. increased gene expression and genome accessibility at associated open chromatin regions or 2. decreased gene expression and genome accessibility at associated open chromatin regions. Negative correlation is defined if 1. decreased gene expression and increased genome accessibility at associated open chromatin regions or 2. increased gene expression and decreased genome accessibility at associated open chromatin regions. All comparisons were made by TP over NP.

|  | **Positively**<br>**Correlated** | **Negatively**<br>**Correlated** |
| --- | --- | --- |
| **Number of dynamic OCRs in association with DEGs** | **2989** | **1660** |
| **Number of DEGs in association with dynamic OCRs** | **1461** | **980** |

**Table 6.** Selected Pathways Associated with Changes of the Open Chromatin Status. IPA Canonical Pathways Analysis.

| Canonical Pathways | -Log(*p*) | Category |
| --- | --- | --- |
| **Signaling by Rho Family GTPases** | 2.80 | Positively Correlated DEGs-OCRs |
| **Actin Cytoskeleton Signaling** | 2.00 | Positively Correlated DEGs-OCRs |
| **NFκB Signaling** | 1.60 | Positively Correlated DEGs-OCRs |
| **Gap Junction Signaling** | 1.66 | Positively Correlated DEGs-OCRs |
| **Glycolysis** | 1.63 | Positively Correlated DEGs-OCRs |
| **Adrenomedullin signaling pathway** | 2.42 | Negatively Correlated DEGs-OCRs |
| **eNOS Signaling** | 2.06 | Negatively Correlated DEGs-OCRs |
| **Relaxin Signaling** | 1.79 | Negatively Correlated DEGs-OCRs |
| **PDGF Signaling** | 1.98 | Negatively Correlated DEGs-OCRs |
| **HIPPO signaling** | 1.68 | Negatively Correlated DEGs-OCRs |

The ATACseq data also allows assessment of transcription factors’ footprints in the open chromatin regions. The bivariate genomic footprinting (BaGFoot) algorithm enables identification of motifs that are located at the deepened protein-binding footprints (Δfootprint depth) sites within the open chromatin regions of increasing DNA accessibility (Δflanking accessibility) from one state to another [37]. The NP and TP myometrial tissues exhibit distinct sets of enriched motifs (Figure S1 and Table S10). The corresponding transcription factors that may occupy at these motifs and are also expressed ALX1, ARID3A, FOXC1, FOXJ2, FOXJ3, HLF, HMGA1, HOXA13, HOXA5, HOXD9, NFIL3, NKX3.1, PBX1, POU6F1, PRRX2, REL and TBP for the TP myometrium. In contrast, the expressing transcription regulators that may bind at enriched motifs in the NP myometrium include EGR1, MDB2, MECP2, NRF1, SP1, SP2, SP3, SP4, ZBTB7B, ZBTB4 and ZNF219. Our data suggest that the interactions between these transcription factors and their genomic target sites in the myometrium are likely subject to the change of chromatin structure at the epigenomic level transitioning in between the two states of myometrium.

### PGR Genome Occupancy in the Myometrial Tissues

Since PGR plays a vital role in mediating the progesterone signaling and enrichment of its DNA motif is found in the open chromatin regions of the myometrium, we next examine the myometrial PGR genome occupancy profiles at both NP and TP stages. Myometrial biopsies from two individual human subjects at each stage were subject to the PGR ChIP-seq assay. Peak calling using data within each individual sample identifies 24095, 9778, 10717 and 8053 PGR occupying intervals in NP1, NP2, TP1 and TP2 myometrial specimens, respectively, after normalization to the same sequencing depth at 9 million reads per sample (Table S11). These PGR binding regions presented in each stage account for 48.3% and 15.0% of open chromatin regions in the NP and TP myometrium, respectively. PGR occupancy is enriched in promoters and 5’ untranslated regions (UTR) in all four myometrial specimens at the expense of the distal intergenic areas (Table 7), implicating a close interaction between PGR and the transcription machinery around the promoters *in vivo* for regulation of gene expression. Notably, the enrichment of myometrial PGR occupancy around the promoter is most prominent in the nonpregnant state (34.3% for NP1 and 35.4% for NP2). On the other hand, such enrichment is relatively lower at the TP state (26.4% for TP1 and 12.0% for TP2) with comparatively higher percentage of PGR binding in the distal intergenic region (Table 7). This finding indicates that re-distribution of PGR genome occupancy occurs concurrently with myometrial remodeling and transcriptomic changes in adaptation to pregnancy.

**Table 7.** Genome-wide PGR occupying sites relative to gene bodies. PGR binding intervals were annotated by the Cis-regulatory Element Annotation System. The promoter is defined as a region within 3 kb upstream of the transcription start site. The downstream of gene body refers to an interval within 3 kb downstream of the transcription termination site.

|  | NP1 | NP2 | TP1 | TP2 | Background |
| --- | --- | --- | --- | --- | --- |
| **Promoter** | **23.6 %** | **25.4 %** | **19.3 %** | **9.6 %** | **2.4 %** |
| **5' UTR** | **10.7 %** | **10.0 %** | **7.1 %** | **2.4 %** | **0.4 %** |
| **Coding Exon** | **4.2 %** | **2.8 %** | **2.1 %** | **0.8 %** | **1.9 %** |
| **Intron** | **33.5 %** | **31.6 %** | **36.2 %** | **41.5 %** | **41.6 %** |
| **3' UTR** | **1.4 %** | **1.5 %** | **1.3 %** | **1.2 %** | **1.4 %** |
| **Downstream of gene body** | **3 %** | **2.2 %** | **2.3 %** | **2.4 %** | **2.2 %** |
| **Distal intergenic region** | **23.4 %** | **26.5 %** | **31.8 %** | **42.0 %** | **50 %** |

The DNA sequences of PGR occupying regions exhibited a significant enrichment of the consensus PGR binding motif [57] in all four myometrial specimens in comparison with the background genome (enrichment *p* value 1 × 10^−18^ [NP1], 1 × 10^−686^ [NP2], 1 × 10^−1105^ [TP1], 1 × 10^−1059^ [TP2], Table S12), while the PGR consensus motif can be found more frequently in the TP than the NP myometrial PGR binding intervals (21.33% [TP1] and 26.20% [TP2] vs. 5.01% [NP1] and 18.57% [NP2], Table S12). Over-representation of binding motifs for transcription factors that have known indications in the myometrium is also seen across all myometrial specimens, including NR3C1 (glucocorticoid receptor) [6, 58], ATF3 [59], FOSL1 [60], JUN [60], KLF9 [61, 62], NF*k*B [63, 64] and STAT5 [54] (Table S12). Similarly, DNA binding motifs for the PGR co-factor FOSL2 [27, 57], smooth muscle differentiation regulator TCF21 [65] and smooth muscle master regulator SRF (CArG)[66, 67] are over-represented throughout the samples, too (Table S12). These data collectively show that the PGR may interact with DNA via both consensus motif dependent and independent manners and such interaction is dynamic during myometrial remodeling. Moreover, our *in vivo* observations provide clinical relevance to previous *in vitro* findings that PGR interacts with other know myometrial and muscle transcription factors in close proximity at cis-acting elements for regulation of myometrial transcriptome.

Further examination of the motif enrichment data identifies candidate myometrial PGR-partnering transcription factors, however, their functions in the myometrium await investigation. The TEAD proteins are known to mediate the Hippo pathway for regulation of muscle fiber size [68] and to interact with AP-1 factors for the smooth muscle fate decision [69]. The motif for TEAD transcription factors is over-represented in PGR binding intervals across all four specimens (Table S12) and expression of TEAD1, TEAD2, TEAD3 and TEAD4 is detected in both NP and TP myometrial tissues (Table S2). The BATF family transcription factors function as a pioneer factor that controls chromatin accessibility [70] and serve as co-regulators for PGR, NR3C1 and AP-1 [71–73]. The binding motif for BATF, members of the AP-1 superfamily, is among the top enriched motifs within PGR binding regions in all four myometrial samples (Table S12) and BATF, BATF2 and BATF3 are all expressed in both stages of the examined myometrium (Table S2). YY1, regulator of chromatin conformation and DNA looping [56] exhibits adequate mRNA levels between NP and TP stages (Table S2). Interestingly, the YY1 binding motif is over-represented surrounding the PGR binding sites of NP myometrial samples but becomes less enriched over background in the TP specimens (enrichment *p* value 1 × 10^−55^ [NP1], 1 × 10^−61^ [NP2], 1 × 10^−20^ [TP1], > 0.01 [TP2], Table S12). These findings implicate potential interactions of PGR with the Hippo signaling pathway, inflammatory response and chromatin conformation regulators for transcription control *in vivo*. Furthermore, the example of stage-associated differential enrichment of the YY1 motif in the PGR binding intervals suggest that PGR may work with various sets of transcription factors to confer phenotype-specific transcriptome profiles.

Next, we determine common and distinct PGR binding intervals between the NP and TP myometrial samples. Union PGR binding peaks across samples were generated based on all PGR binding intervals in four specimens to document the presence and absence of each PGR binding site in individual specimens. Total 30613 union PGR binding peaks are found (Figure 5 and Table S13). The numbers of union peaks present in each specimen are 24019, 9491, 10402 and 7928 for NP1, NP2, TP1 and TP3, respectively (Figure 5). NP1 and NP2 myometrial tissues share 7323 union peaks while that number between TP1 and TP2 is 4839 (Figure 5). Among these 7323 peaks in the NP myometrium, binding motifs of PGR, STAT transcription factors, SRF, MyoD and SMAD2 are over-represented (Figure 6 and Table S14). Enrichment of these motifs are also found among the 4839 PGR occupying sites in the TP myometrium (Figure 6 and Table S14). Consistent with motif analysis results on individual specimens (Table S12), the PGR consensus motif exhibits higher levels of enrichment in the PGR occupying intervals of the TP myometrium in comparison with that of the NP stage (Figure 6). In addition, STAT, MyoD and SMAD2 motifs also show increased enrichment in the TP myometrium while the CArG motif has stronger over-representation in NP specimens (Figure 6 and Table S14). These observations indicate adjustment of partnership with other transcription factors at different stages is a common process among specimens beyond individual variations.

**Figure 5.**
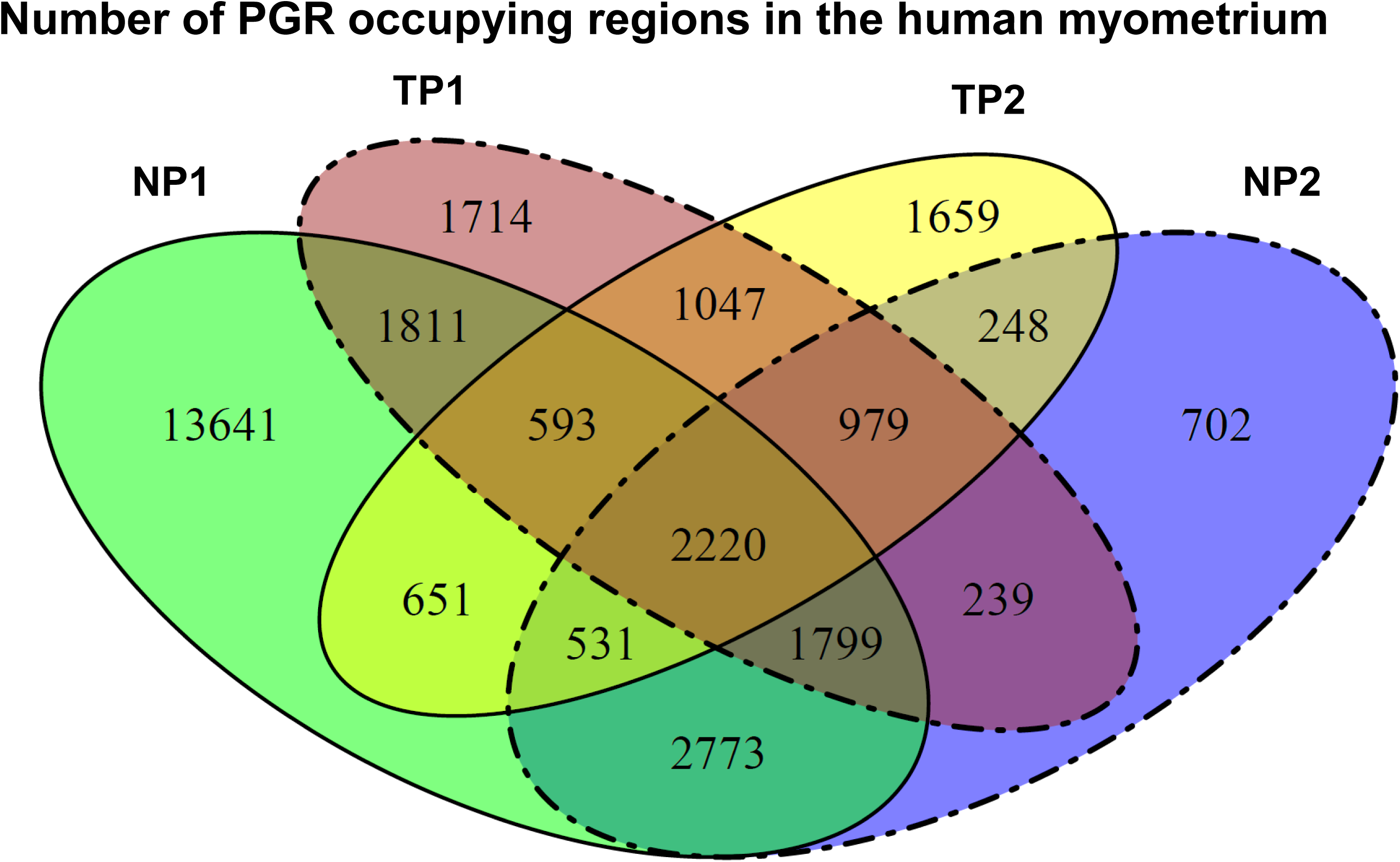
Number of PGR occupying regions in the genome of the human myometrium. PGR ChIP-seq was performed in two separate specimens at either NP or TP stages. The flower Venn diagram illustrates the numbers of overlapping and distinct PGR occupying sites among the four specimens.

**Figure 6.**
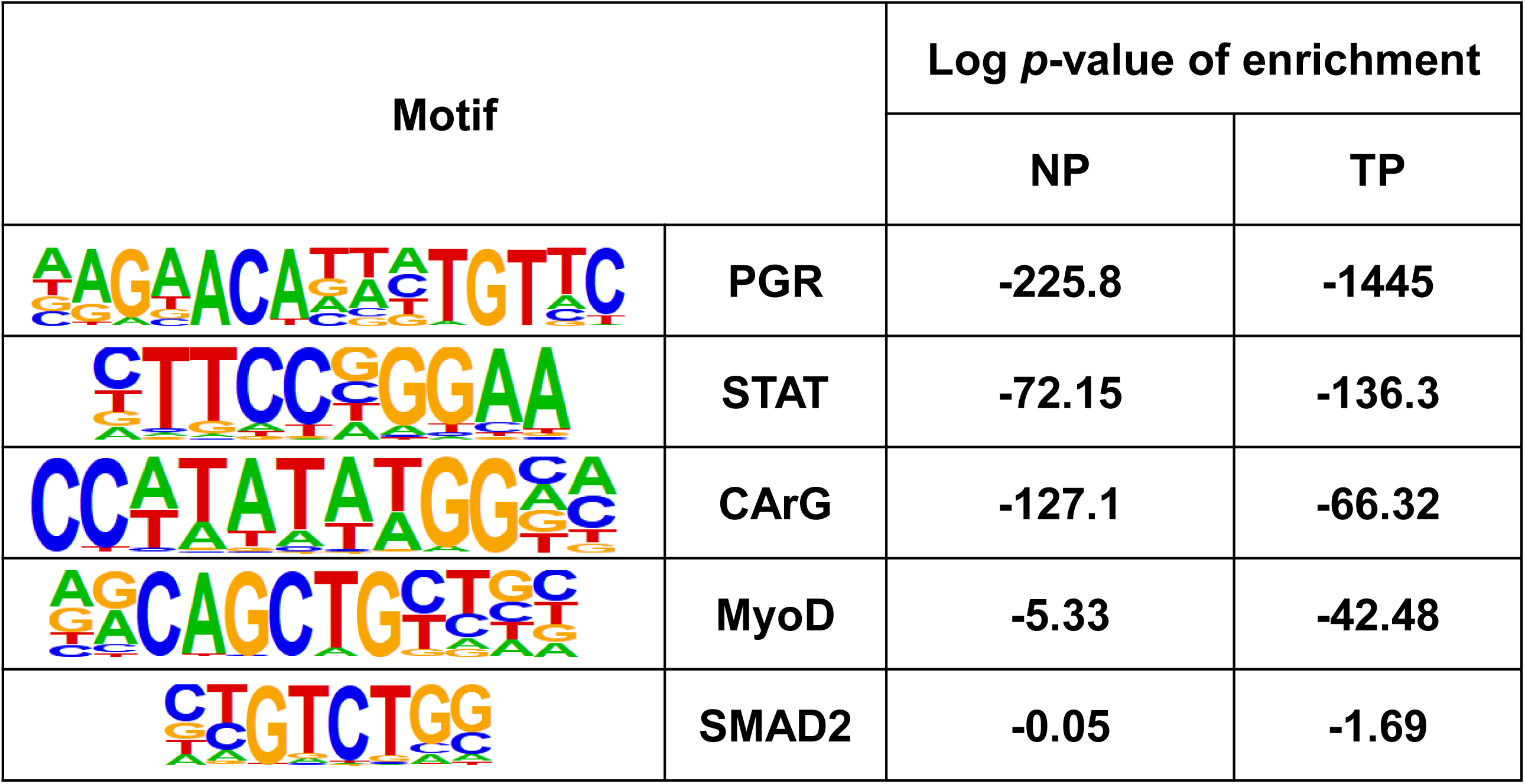
Selected Enriched motifs in common PGR-binding intervals at each stage.

Further examining the stage-associated PGR occupying regions identified 2220 genomic regions that are shared between the 7323 and 4839 PGR binding intervals while 2773 and 1047 have distinct presence in NP and TP myometrium, respectively (Figure 5). In addition to differences in genomic locations, the compositions of transcription factor binding motifs also show divergent profiles among these three groups of PGR occupying regions (Table S15). Among the 95 motifs that are compiled from the top 50 enriched motifs of each of the three groups, the enrichment p values reveal a major difference on the extent of these motifs’ presentation between the NP-distinct and TP-distinct PGR occupying regions (Figure 7). The TP distinct regions are highly enriched for the consensus binding motifs for progesterone and glucocorticoid receptors, AP-1 proteins and TEAD transcription factors, in contrast to the relatively lower enrichment in the NP distinct regions. On the contrary, NP distinct sites show strong over-representations on motifs of ETS transcription factors, nuclear transcription factor Y, CTCF, YY1, Sp1 and NRF1, which are much less enriched in the TP distinct regions. Notably, the 2220 shared regions have a motif enrichment profile similar to that of TP rather than NP distinct regions (Figure 7). This suggests PGR at the NP stage may interact with a wider spectrum of transcription regulators while it likely works with a more focused set of transcription factors in the TP myometrium. The data collectively suggest that the dynamics of PGR binding locations between the two states of myometrium allow various combinations of transcription factor binding motifs available for corresponding regulators to interact with PGR at a state-specific manner.

**Figure 7.**
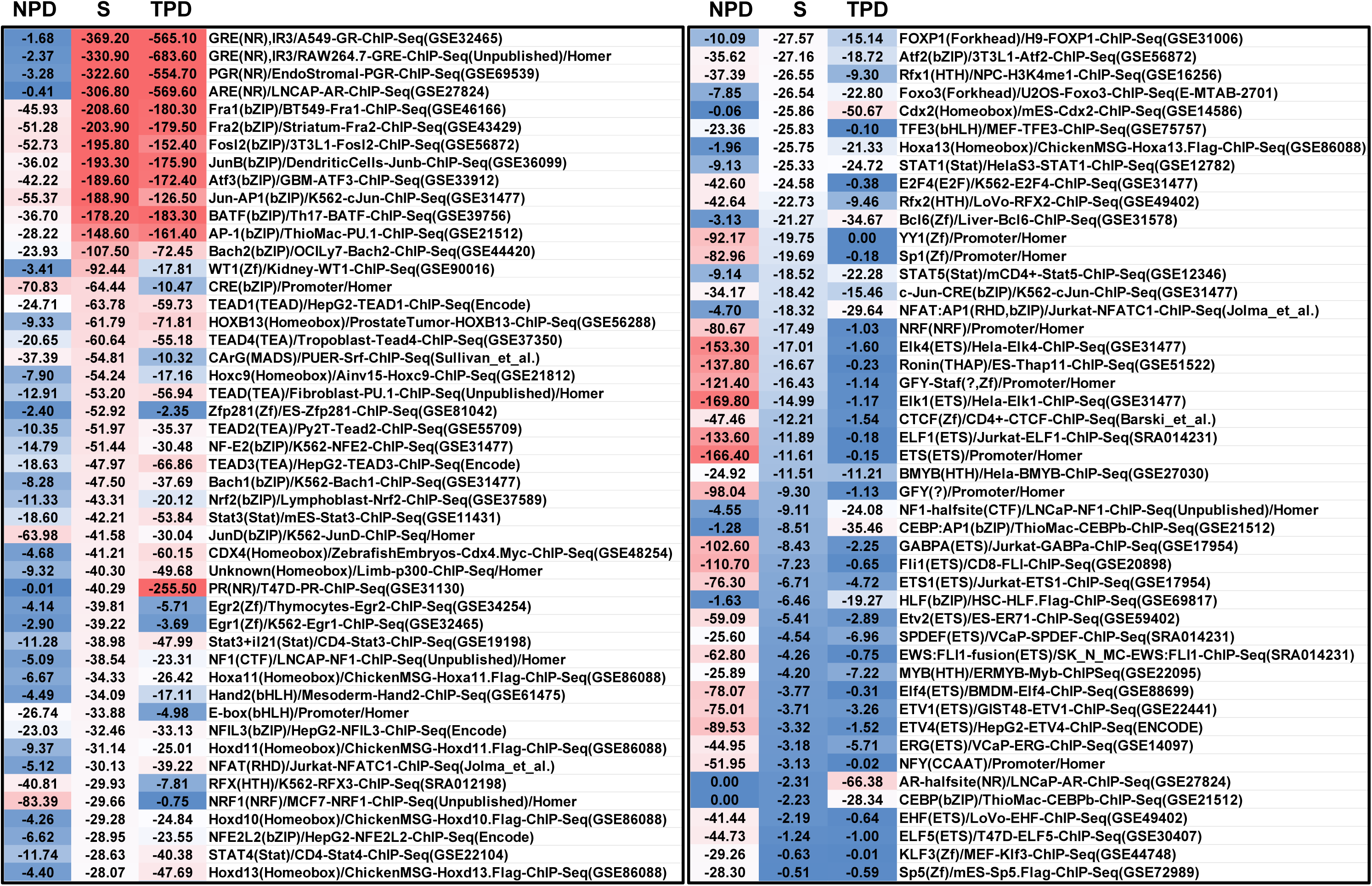
Top enriched transcription factor binding motifs in 2773 NP distinct (NPD), 1047 TP distinct (TPD) and 2220 NP-TP shared (S) PGR occupying sites. Top 50 motifs from each group are compiled and displayed with log p values. Color codes are depicted with the red-white-blue pattern with red showing the strongest enrichment and blue being the least enriched.

### Candidate PGR Direct Downstream Target Genes in the Myometrium

The association between the PGR genome occupancy and the myometrial transcriptome is assessed by identifying the nearest genes to 2220 shared and 2773 and 1047 distinct PGR binding intervals between the NP and TP myometrium. Among these 6040 PGR occupying regions, 4355 (72%) are associated with 2203 myometrial active genes (Table 8 and Table S16). These 2203 candidate PGR direct downstream target genes can be further sub-grouped into 1431 NP-distinct, 183 TP-distinct and 589 NP-TP shared active genes based on their association with distinct or common PGR binding intervals (Table 8), in which Ingenuity Pathway Analysis identified 51, 37 and 166 canonical signaling pathways, respectively, that have enrichment *p* value less than 0.05 (Table S17). The top 50 associated pathways from each of the three groups manifest divergent profiles (Figure 8). Notable example pathways that are selectively enriched in individual groups include “estrogen receptor signaling” and “DNA Methylation and Transcriptional Repression Signaling” for the NP distinct group, “IL-10 Signaling” and “Inhibition of Matrix Metalloproteases” for the TP distinct group, and “Actin Cytoskeleton Signaling” and “p53 Signaling” for the NP-TP shared group (Figure 8 and Table S17). In contrast, no canonical pathway has the enrichment *p* value less than 0.05 across all three groups. These findings suggest that each group of PGR binding regions may contribute to regulation of a cohort of genes that cater to specific functions for physiological adaption on pregnancy.

**Figure 8.**
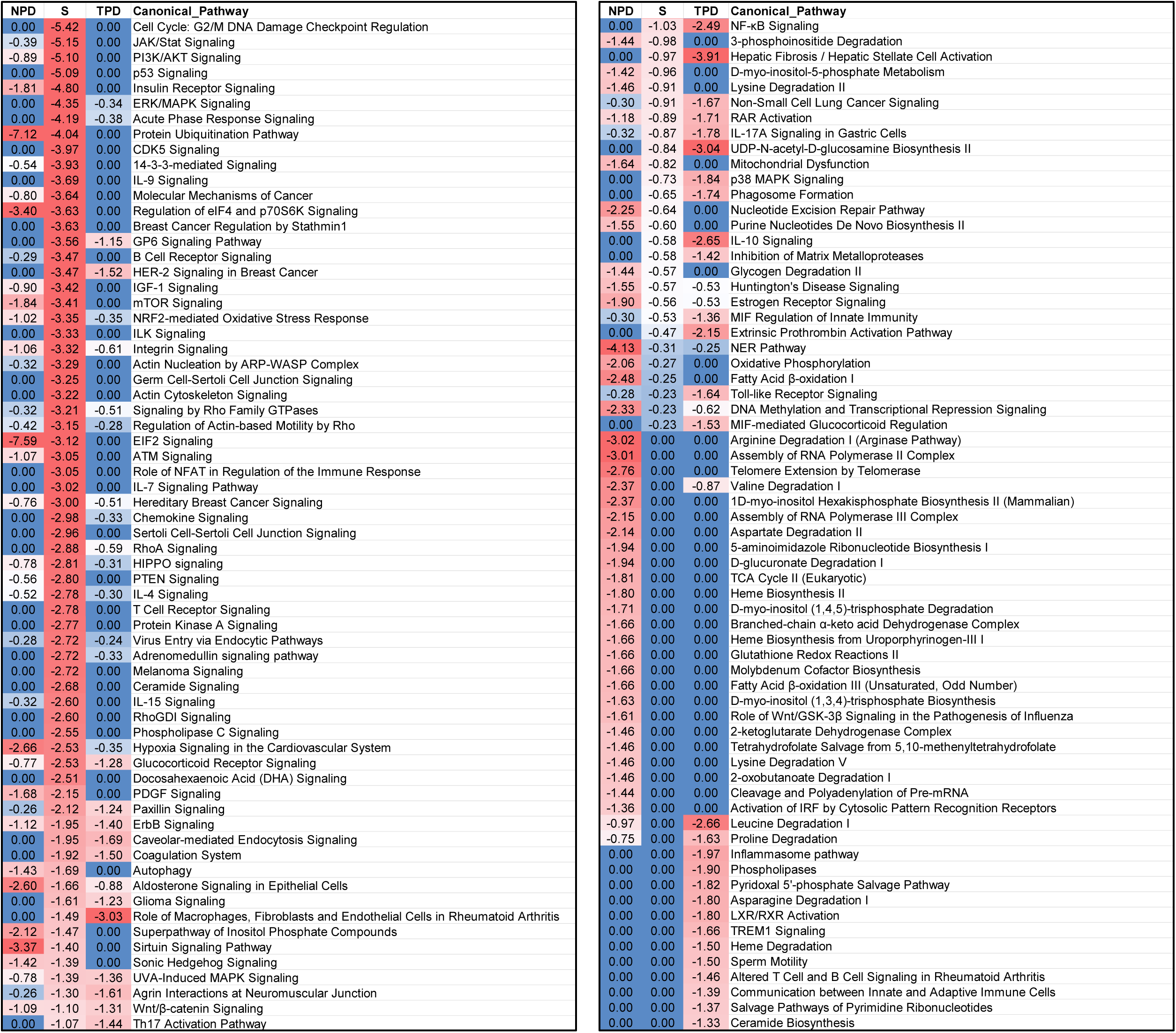
Top enriched canonical pathways in 1431 NP-distinct (NPD), 183 TP-distinct (TPD) and 589 NP-TP shared (S), PGR-linked active genes. Top 50 enriched pathways from the Ingenuity Pathway Analysis from each group are compiled and displayed with log p values. Color codes are depicted with the red-white-blue pattern with red showing the strongest enrichment and blue being the least enriched.

**Table 8.** The association between PGR occupancy and myometrial active genes. A list of nearest genes to the 2220, 7323 and 4839 shared and distinct PGR occupying intervals was identified and further filtered with the criteria of having the maximum FPKM value greater or equal than 1 among the 6 myometrial RNAseq samples to catalog the PGR-associated active genes.

| NP Distinct | Shared by NP and TP | TP Distinct |
| --- | --- | --- |
| <u>PGR occupying intervals examined</u> |  |  |
| 2773 | 2220 | 1047 |
| <u>Number of PGR occupying intervals in association with active genes in the myometrium</u> |  |  |
| 2252 | 1510 | 593 |
| <u>Number of myometrial active genes in association with PGR occupying intervals</u> |  |  |
| 1431 | 589 | 183 |

PGR occupancy on genomic regions in association with genes of interest are further examined in additional myometrial specimens, two each from NP and TP stages (Table 9). The glycolytic genes *ENO1* and *LDHA* display reduced binding events in the TP myometrium compared with that in NP. Given that both genes exhibit higher expression levels in the TP samples (Figure 3), these findings together indicate a negative correlation between PGR occupancy and mRNA levels. PLCL1 functions to attenuate prostaglandin-dependent calcium signaling (US Patent 20160161487A1). *PLCL1* is expressed in human myometrium (Figure 3 and Table S2) [40] and is a PGR downstream target [74]. PGR occupancy in a region 93kb upstream of the *PLCL1* gene can be detected in both cohorts of myometrial specimens (Table 9 and Table S13), which may serve as a functional regulatory element for the myometrial *PLCL1* gene. While only a few genomic regions were tested, the reproducible detection of PGR occupancy in different cohorts of myometrial specimens validates the genome-wide ChIP-seq results. Moreover, these findings also identified candidate cis-acting elements through which PGR may regulate expression of glycolytic genes *ENO1* and *LDHA* as well as the contractility-associated gene *PLCL1*.

**Table 9.** PGR occupancy at selected genomic loci in a second cohort of myometrial specimens. ChIP-qPCR was performed in a separate set of NP (n=2) and TP (n=2) myometrial tissues for PGR occupancy test. Technical triplicates for each qPCR reaction was performed and the mean and the standard deviation of triplicates was used for calculation of PGR binding events. Untr12 located in a gene desert serves as a negative control.

| Binding Events Detected Per 1000 Cells |  |  |  |  |  |
| --- | --- | --- | --- | --- | --- |
| Gene | Region of Interest | NP3 | NP4 | TP3 | TP4 |
| *ENO1* | chr1:8938001-8939564 | 25.93 ± 1.89 | 12.46 ± 1.44 | 0.91 ± 0.38 | 0.71 ± 0.93 |
| *LDHA* | chr11:18415880-18416428 | 30.15 ± 3.49 | 11.42 ± 1.45 | 1.56 ± 0.42 | 2.35 ± 0.68 |
| *PLCL1* | chr2:199107091-199107777 | 119.13 ± 8.45 | 7.77 ± 1.76 | 9.44 ± 0.31 | 6.40 ± 0.79 |
| *Untr12* | Chromosome 12 | 0.15 ± 0.15 | 0.00 ± 0.00 | 0.13 ± 0.23 | 0.00 ± 0.00 |

## Discussion

### Novelty of the study

The present study documents the molecular landscape of the NP and TP myometrium as well as the changes between these two stages. While previous genome-wide studies identify candidate mechanisms for the switch from nonlabor to labor states [40, 75, 76], results presented here report the way the myometrium reorganizes and adapts from the nonpregnant to the term pregnant state. This is exemplified by the uniform increase of mRNA of parturition associated genes *OXTR, GJA1, ZEB1* and *PLCL1* during myometrial remodeling followed by the divergent change of expression patterns (Figure 1). Notably, myometrial *ZEB1* and *PLCL1* mRNA levels both decrease from the term nonlabor to the term labor state [40, 43], which contrasts with our observations that these two genes exhibit an increase of expression from the nonpregnant to the term pregnant state (Figure 1). At the genome-wide level, 60 of 126 genes that have significant changes of mRNA levels between the term nonlabor and the term labor state either show no change (55 genes) or in an opposite direction (5 genes) of expression in our studies [75]. While this discrepancy could result from data variances of different cohorts of human subjects, it is also possible that these 60 genes could be under tight regulation at the nonpregnant, the term nonlabor and the term labor states for potential stage-specific functions. These observations justify future examinations on the timely change of myometrial transcriptome at multiple states for a high-resolution view of gene expression dynamics. Collectively, our transcriptome data broadens the spectrum of the knowledge on myometrial gene expression dynamics and set the foundation for future meta-analysis to understand the genetic program for myometrial remodeling between the nonpregnant and term pregnant states. The molecular pathways revealed here can also serve as a reference for studying genetic and environmental impacts that derail normal pregnancy leading to disorders such as preterm birth and dystocia.

The assessible genome maps offers a blueprint to study genetic and epigenetic regulation of the myometrial transcriptome. The identified open chromatin regions likely house cis-acting elements that participate in regulation of the myometrial transcriptome and they could be subject to epigenetic control. For example, binding motifs of the estrogen receptor and the cell cycle regulator E2F1 are over-represented in the open chromatin regions found only in the NP myometrium (Table 4), which reflects the higher expression levels of genes for transcription and gene control (Table 2) and implicates the preparation of smooth muscle proliferation for the early gestation myometrial expansion under the control of estrogen receptor. Another example is the parturition associated gene MEF2A [51]. Enrichment of the MEF2A binding motif in the class III open chromatin regions (Table 4), suggest that accessibility of MEF2A to its target sites could be another layer of regulation of its action in addition to changes in expression levels [51]. Our genome-wide integrative analysis of genome accessibility and transcriptome have identified potential regulatory elements and associated molecular pathways that may be affected the chromatin accessibility (Table S8 and table S9). These findings pave the road for future in depth study of the epigenetic impact on myometrial transcriptome changes.

Given that progesterone is the major steroid hormone for pregnancy and has a clinical indication on preventing premature labor, the focused genome-wide myometrial PGR occupancy study provides a map to identify PGR-associated cis-acting elements and their candidate downstream target genes for further investigation on the mechanism of action of PGR. Notably, the impact of losing uterine p53 on gestation length has been reported in the mouse model [49]. In our study, the “p53 Signaling” is among the top enriched pathways in the putative PGR direct downstream targets (Table S17). This finding suggests a potential interaction between PGR and p53 signaling in the myometrium. Meanwhile, the over-represented transcription factor binding motifs surrounding the PGR occupying sites suggest candidate PGR interacting partners for myometrial gene regulation. For instance, the AP-1 proteins are known to partner with PGR for gene regulation as exemplified by a few parturition maker genes [27]. We found that binding motifs of many AP-1 subunits are among the top enriched motifs in the myometrial PGR occupying regions (Table S14 and Figure 7), suggesting a potential genome-wide partnership between PGR and AP-1. Our results not only provide human relevance to the known signaling pathways at the *in vivo* level, but also allow identification of novel targets, either working with or under control of PGR, for future research.

### An Indication of Progesterone Receptor on Myometrial Metabolism and Remodeling

The elevated expression of genes for the glycolytic pathway and lactate production in the term pregnant myometrial specimen in comparison to the nonpregnant state have two physiological implications. First is the lactate production and the myometrial contractility. Myometrial lactic acidosis is observed in women who have dysfunctional labor [77]. It has been shown that acidic pH abrogates myometrial contraction [78]. Lactate treatment can reduce spontaneous contraction [79] and suppresses uterine inflammation and preterm birth [80]. Lactate is converted from the glycolysis product pyruvate in the muscle system by the lactate dehydrogenase. Decreased muscle *Ldha* expression, which encodes a subunit of the lactate dehydrogenase, leads to a reduction of pyruvate-to-lactate conversion [81]. Our transcriptomic data indicates that the pathway for glycolysis and lactate production is set for an increase of processing capacity at the term state compared to that in the nonpregnant myometrium (Figure 3 and Table S2). Moreover, higher myometrial utilization of glucose at term pregnant state is also supported by the change of expression pattern of glucose responsive genes (Table S4 and Table S5). Our findings suggest that the myometrial transcriptome adapts to ramp up the lactate production capacity, which contributes to maintaining the myometrial quiescence. The second note is on the energy metabolism and molecular biosynthesis. Our transcriptomic data shows that, in the TP myometrium, an increase of HIF1A molecular activity likely occurs based on the expression pattern of HIF1A downstream targets (Table S4 and Table S5). The rise of HIF1A activity suggests a hypoxic myometrium at term. Under the anaerobic condition, increased glycolysis can generate ATP to compensate the need of energy for maintaining cellular functions. On the other hand, at normoxia, the increased capacity for higher flux of glycolysis can supply the intermediates for amino acid synthesis to generate building blocks for myometrial expansion. The increased glycolytic gene expression in response to pregnancy would fit these hypotheses for the change of myometrial functions (Figure 3). Lastly, the binding of PGR in the genomic loci of ENO1 and LDHA links PGR to the regulation of myometrial metabolism (Table 9), which awaits further investigation.

### Limitation and future direction

The resolution of molecular profiling data, integration of multiomics information and small sample size on the ChIPseq, RNAseq and ATACseq assays are major limitations for the present study. The genome-wide data here is a summary observation of many cells that reside in pieces of myometrial tissues. While the smooth muscle cells make up the majority of the mass, readings from other cell type such as blood and vascular cells are also incorporated as part of the summary results, which may reduce the signal-to-noise ratio of the information on the cell type of interest. Despite potential contribution of other cell type transcripts, the myometrial transcriptome data exhibits a strong association with known smooth muscle and myometrial programs (Table 2 and Table 3), the ATACseq data establishes close links between putative cis-acting elements and signaling pathways for muscle functions (Table 6), and the PGR cistrome data detects enrichment of binding motifs for smooth muscle master regulators SRF and MyoD (Figure 6). These examples demonstrate that the current data can provide significant information to support existing knowledge and supply novel findings. The emerging molecular profiling assays at the single cell level would provide a greater resolution than current results obtained with more established technology, leading to further dissection of each cell’s role and the underlying molecular events in a given myometrial state.

Another limitation on the data resolution is that the antibody used for generating the current PGR ChIPseq results detects both major PGR isoforms in the myometrium, the PGR-A and PGR-B. These two PGR isoforms differ in the AF-3 transactivation domain that only exists in the PGR-B. Both isoforms are functionally capable of modulating transcription of common and distinct sets of downstream genes depending on the context [25]. Therefore, the stoichiometric ratio between the two isoforms in different stages of myometrium needs to be factored into interpretation of the effect of PGR on expression of associated genes. For studies on human tissues, there is an urgent need to develop the isoform-specific antibodies for PGR, perhaps through targeting the protein domains with isoform-specific conformations.

With respect to the limitation on integration of multiomics information, in this manuscript the PGR occupancy and myometrial transcriptome association study is based on a stringent criterion that only link one nearest gene to each individual PGR occupying site (Table 8). The rationale to use such as stringent criterion is based on the observation that majority of the PGR occupying sites are located near gene bodies (Table 7). PGR occupied cis-acting elements likely impose a greater influence on the myometrial transcriptome according to emerging transcription control models that each gene is subject to regulation by multiple enhancers and vice versa in a topological associated domain formed via chromatin conformation [82, 83]. Future integration of chromatin conformation data, generated from the high throughput chromatin conformation assay, would better assign PGR binding cis-acting elements with associated genes, especially on the long-distance interactions, to provide correlation information closer to the true molecular events.

The small sample size on ATACseq and PGR ChIPseq, in part, also accounts for limitation of the present study. The size of biopsied specimens and the required amount of tissues for the chemistry of these assays weight heavily on the sample size for the study, especially for ATACseq and ChIPseq assays. Moreover, the content and texture of the muscle tissue render challenges on the current chemistry of chromatin immunoprecipitation for ChIPseq and library preparation for ATACseq. These factors jointly hinder our ability to examine samples in a larger cohort at the genome-wide scale. Emerging novel technologies including CUT&Tag, low cell input protocols for ChIPseq and ATACseq, and using isolated nuclei from frozen tissues as containers of chemistry could overcome some of the technical obstacles and allow future examination on larger cohorts of specimens [84, 85].

Our data document the molecular profiles at two ends of the physiological spectrum of myometrial tissues. Aside from the different functional and structural states, the nonpregnant specimens were at a low serum progesterone environment while the term pregnant samples were exposed to high levels of serum progesterone. Therefore, confounding factors from the physiological states of the muscle tissues and the differential levels of serum progesterone levels shall be considered at interpretation of the molecular changes between the nonpregnant myometrium at the proliferative phase of the menstrual cycle and the term pregnant myometrium. Nevertheless, our findings provide the baseline information that can be integrated with observations from *ex vivo* explant culture or *in vitro* cell culture models to tease out underlying mechanisms that regulate myometrial remodeling. The large differences on the molecular profiles between these two states also suggest that future examinations of myometrial tissues on stages earlier than the term pregnancy may provide a better resolution in pregnancy dependent molecular dynamics of the myometrium, should the samples could be obtained at a sufficient amount for experimentation.

## Non-Standard Abbreviation

AP-1: Activator protein 1
ATACseq: Assay for Transposase-Accessible Chromatin using sequencing
BaGFoot: Bivariate genomic footprinting
cAMP: Cyclic adenosine monophosphate
ChIP-qPCR: Chromatin Immunoprecipitation quantitaitve polymerase chain reaction
ChIPseq: Chromatin Immunoprecipitation Sequencing
DAVID: The Database for Annotation, Visualization and Integrated Discovery
DEG: Differentially expressed gene
eNOS: Endothelial nitric oxide synthase
ERK: Extracellular signal-regulated kinases
FDA: United States Food and Drug Administration
FPKM: Fragments per kilobase of exon per million fragments
GTPase: Guanosine triphosphate hydrolase
HOMER: Hypergeometric Optimization of Motif EnRichment
IPA: Ingenuity Pathway Analysis
MACS2: Model-based Analysis of ChIP
NFAT: Nuclear factor of activated T-cells
NFkB: Nuclear Factor Kappa B
NP: Nonpregnant
OCR: Open chromatin regions
PDGF: Platelet-derived growth factors
PI3K: Phosphoinositide 3-kinase
STAT: Signal transducer and activator of transcription
TEAD: TEA Domain Transcription Factor
TNF: Tumor necrosis factor
TP: Term Pregnant
WNT: Wingless-type MMTV integration site family

## Acknowledgement

This work was supported by the Intramural Research Program of the National Institutes of Health: Project Z1AES103311-01 and March of Dime. We thank the NIEHS Epigenomics and DNA Sequencing Core Laboratory for the technical assistance and Ms. Sylvia Hewitt for critical comments.

## Author Contribution

SPW and FJD designed and wrote the manuscript. MLA obtained permission from the Institutional Review Board to perform all work, collected the patient consents and myometrial specimens and wrote the corresponding section of the Methods section. SPW, OME and XL performed experiments and associated data analysis. TW, LZ and SPW conducted bioinformatic analyses. TW and LZ wrote the bioinformatic part of the Methods section.

## Supplemental Information

### Legends of Supplemental Figures and Tables

**Figure S1.**
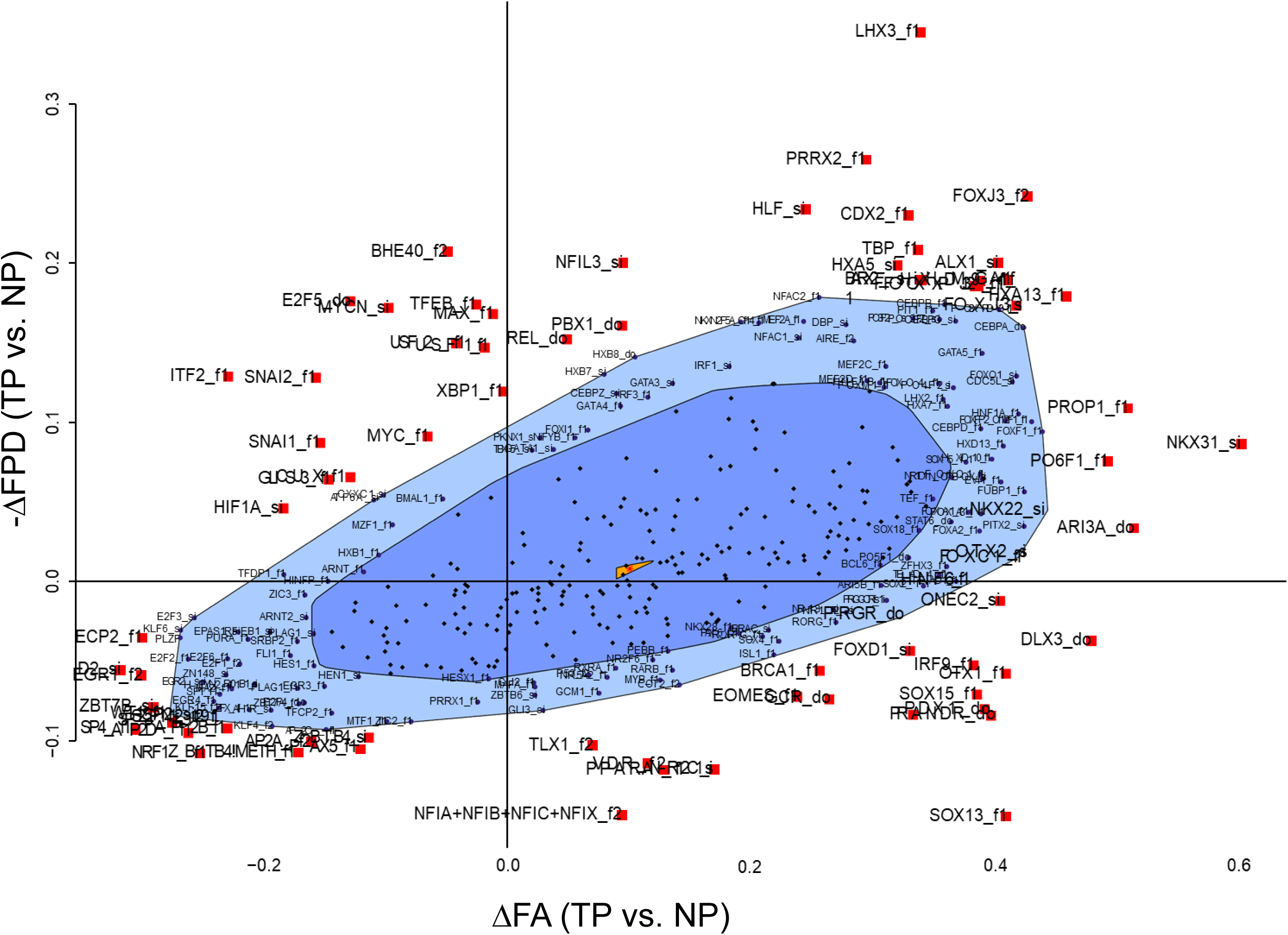
Bag plot depicting change of DNA accessibility (ΔFA) and alteration in the depth of protein-binding footprints (ΔFPD) from the NP to the TP stages. Motifs with statistical significance are labeled in red squares.

**Table S1.** List of human myometrial specimens.

**Table S2.** Active Genes in Human Myometrium

**Table S3.** A complete list of Gene Ontology analysis of Table 2.

**Table S4.** Predicted molecular activities by Ingenuity Pathway Analysis Differentially Expressed Genes between NP and TP human myometrium.

**Table S5.** Predicted molecular activities by Ingenuity Pathway Analysis on Differentially Expressed Genes between 18.5dpc and virgin mouse myometrium (GSE17021).

**Table S6.** Genome coordinates of open chromatin regions in human myometrial specimens. “1” and “0” represent presence and absence, respectively, in the NP and TP myometrium. Classes are defined in Figure 4.

**Table S7.** Motif enrichment analysis by HOMER known motif in the three classes of open chromatin regions that are defined in Figure 4.

**Table S8.** Motif enrichment analysis on the DEG-associated dynamic OCRs.

**Table S9.** Canonical Pathways that are linked to dynamic OCRs associated DEGs.

**Table S10.** Transcription footprinting assessment in the NP and TP myometrial specimens by BaGFoot.

**Table S11.** Genome coordinates (Hg19) of PGR occupying intervals in myometrial tissues of four human subjects.

**Table S12.** Enriched motifs in each individual myometrial specimen. Data is from the output files of the HOMER unkown motif search algorithm.

**Table S13.** Union peak information of PGR binding intervals across all four myometrial specimens. In the “UnionPeak-vs-Sample” section, 1 and 0 denote presence and absence of the PGR binding peak in each individual specimen.

**Table S14.** Enrichment analysis on 428 transcription factor binding motifs in common PGR occupying sites within each stage of myometrial specimens.

**Table S15.** Enrichment analysis on 428 known transcription factor binding motifs in shared and distinct PGR occupying regions between NP and TP groups.

**Table S16.** Association between PGR occupancy and myometrial active genes.

**Table S17.** Canonical pathways in association with PGR occupancy and myometrial active genes. 1431, 589 and 183 active genes that are linked to NP-distinct, NP-TP shared and TP-distinct PGR occupying sites, as depicted in Table 8, were subject to Ingenuity Pathway Analysis. Pathways that have enrichment *p* value less than 0.05 are marked light blue.

